# PhoCoil: An Injectable and Photodegradable Single-component Recombinant Protein Hydrogel for Localized Therapeutic Cell Delivery

**DOI:** 10.1101/2024.05.07.592971

**Authors:** Nicole E. Gregorio, Cole A. DeForest

## Abstract

Hydrogel biomaterials offer great promise for 3D cell culture and therapeutic delivery. Despite many successes, challenges persist in that gels formed from natural proteins are only marginally tunable while those derived from synthetic polymers lack intrinsic bioinstructivity. Towards the creation of biomaterials with both excellent biocompatibility and customizability, recombinant protein-based hydrogels have emerged as molecularly defined and user-programmable platforms that mimic the proteinaceous nature of the extracellular matrix. Here, we introduce PhoCoil, a dynamically tunable recombinant hydrogel formed from a single protein component with unique multi-stimuli responsiveness. Physical crosslinking through coiled-coil interactions promotes rapid shear-thinning and self-healing behavior, rendering the gel injectable, while an included photodegradable motif affords on-demand network dissolution via visible light. PhoCoil gel photodegradation can be spatiotemporally and lithographically controlled in a dose-dependent manner, through complex tissue, and without harm to encapsulated cells. We anticipate that PhoCoil will enable new applications in tissue engineering and regenerative medicine.

## Introduction

Hydrogels provide specialized material environments that are conducive to studying cells in 3D as extracellular matrix mimics and delivering therapeutics in a controlled manner (*1–3*). However, many of the most popular hydrogels used in biological or biomedical applications suffer from either a lack of tunability or shortcomings in their biocompatibility or biodegradability. In the case of naturally derived materials consisting of biomolecules extracted from tissues, precise control over material properties remains elusive or requires additional synthetic modifications (*4*, *5*). Moreover, batch-to-batch variability in these natural materials can be an insurmountable hurdle to both experimental consistency and approval for clinical use (*6*, *7*). While biomaterials research has also favored synthetic polymer-based materials for their ability to be readily modified and controlled by the user, these materials are infrequently chosen for therapeutic applications due to concerns of low cytocompatibility and biodegradability or high immunogenicity, as well as their intrinsic polydispersity (*5*, *7*, *8*). On top of this, these materials often require significant expertise in synthetic chemistry to install the reactive and responsive chemical handles that enable user-specified network customization, rendering them unreachable by many biologically focused groups.

Over the past 25 years, recombinant protein-based hydrogels have become an enticing alternative to natural and synthetic polymer hydrogels. Recombinant protein hydrogels can bridge the gap – retaining both a highly biocompatible and bioresorbable makeup similar to natural protein-based gels, while providing greater user-defined tunability without the need for synthetic chemistry (*5*, *7*, *8*). The ability to control the mechanical and responsive characteristics of these protein networks is genetically encoded, arising from user-defined variations at the amino acid and fusion protein levels, which can be readily modified through well-established cloning techniques already common across the biological sciences. Furthermore, recombinant protein production can be accomplished with relatively simple and widely utilized bacterial protein expression techniques and common laboratory equipment. Recombinant protein-based materials have the added benefit of being familiar substrates for cells; they can easily be designed to promote cell-material interactions through the inclusion of cell binding motifs and can be processed and broken down by pre-existing cellular machinery (*5*, *8*). Collectively, these properties make recombinant protein hydrogels well-suited towards biomedical applications such as localized therapeutic cell delivery.

In this work, we sought to design a recombinant biomaterial system that would enable minimally invasive cell delivery and user-controlled cell release. To achieve this, we envisioned a material that was both injectable and photodegradable, while retaining high biocompatibility in its intact and degraded states. Our group recently developed an injectable recombinant protein-based hydrogel that utilizes coil motifs as physically crosslinking domains linked together via an unstructured flexible linker called XTEN (*9*). The shear-thinning and self-healing nature of this telechelic, single-component hydrogel maintained high viability of encapsulated cells throughout and following injection both *in vitro* and *in vivo*. While this physically crosslinked XTEN hydrogel slowly degrades through surface erosion, the timing of degradation is beyond user control.

Building on this work, we sought to introduce photodegradability to the material to enable user-actuated spatiotemporal control of network degradation. We anticipated that the insertion of a previously evolved protein called PhoCl (*10*) at the center of the XTEN linker would introduce this photodegradable response (**Figure 1**). PhoCl is a photocleavable protein whose peptide backbone undergoes irreversible cleavage at the chromophore in response to 405 nm light, providing both a cytocompatible and highly bioorthogonal trigger for 4D control of protein cleavage (*10–12*). Previous work from our group and others has shown PhoCl’s utility in introducing photoresponsivity to hydrogels (*13–15*). The resulting material, referred to as “PhoCoil”, achieves the intended combination of chemical, physical, and stimuli-responsive properties (biocompatible, injectable, photodegradable) which are all directly specified in its molecular design. Our PhoCoil material offers unique benefits over previously reported photodegradable recombinant protein-based materials (*16–23*) as it is formed directly from a single protein component, requires no small molecules to initiate gelation and/or photocleavage, is stable under ambient light for easy handling, is readily injectable in its gel state, and degrades in response to cytocompatible visible light. In this work, we thoroughly characterize PhoCoil’s physical and light-responsive properties as well as provide proof-of-concept studies highlighting its potential as an injectable cell delivery and triggered release platform.

**Fig. 1.**
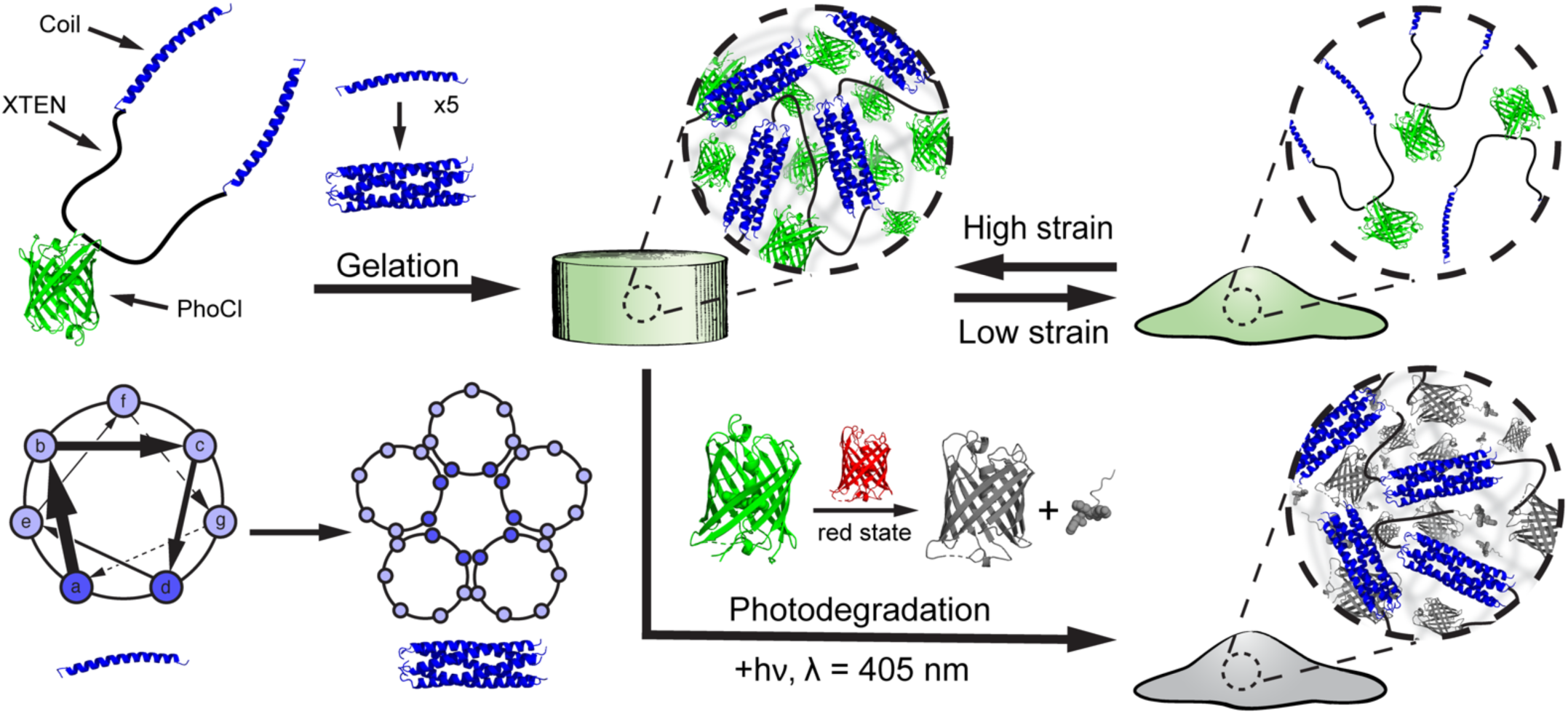
PhoCoil structure and stimuli-responsive properties. PhoCoil hydrogels are physically self-assembled through homopentameric coiled-coil bundle formation. These bundles are held together by noncovalent interactions along the length of the helices, primarily between amino acids at the a and d positions.The noncovalent nature of these associations endows the bulk gel with shear-thinning and self-healing properties in response to applied force. The network degrades in response to 405 nm light due to the cleavage of the peptide backbone within the chromophore of PhoCl. During this process, PhoCl temporarily occupies a red state prior to the diffusive separation of the two protein halves and subsequent network disassociation. Protein Data Base ID Codes: PhoCl intact (green state) – 7DMX, PhoCl cleaved (red state) – 7DNA, PhoCl cleaved and dissociated (colorless state) – 7DNB, Coiled-coil - 1MZ9.

## Results

### Protein design

The PhoCoil protein represents a new design that builds on our team’s prior efforts in the recombinant protein hydrogel space (*9*) by encoding photodegradability into the primary protein sequence. PhoCoil’s design is similar to that of a symmetric pentablock copolymer containing three distinct segments arranged in an ABCBA pattern that each individually contribute to its macromolecular properties (**Figure 1**). This telechelic protein has self-associating coil domains at both termini (Block A), allowing for network formation through physical interactions between individual protein units. These flanking coils are naturally derived from the cartilage oligomeric matrix protein sequence and form homopentameric bundles that associate through noncovalent interactions along the length of the coils (*24*, *25*). Due to these physical interactions, coil association is reversible under varying force. For example, high-strain conditions pull apart the coiled-coil structures, resulting in shear thinning of the network and a transition to a liquid-like state. Subsequent return to a low-strain state allows the coiled coils to reform, returning the network to its original, gel-like state. This shear-thinning and self-healing behavior of the network enables injection through small-gauge needles post gelation. We take advantage of coiled-coil domains, as they have been widely utilized to create physically crosslinked protein biopolymer networks, enabling a variety of applications in injectable, sustained drug delivery (*8*, *26–33*).

At the center of the protein is PhoCl (Block C), a green fluorescent protein that undergoes irreversible cleavage in response to 405 nm light, providing the network with its photodegradable properties (*10*). PhoCl cleavage occurs in the chromophore of the protein, where visible light induces a β-elimination reaction that results in scission of the peptide backbone, producing a short-lived red state of the protein. This red state is temporarily captured in our hydrogels and dissipates as the two halves of the protein diffuse away from each other, resulting in the dissolution of the gel network as the coiled-coil bundles are no longer covalently linked. Given this cleavage mechanism, PhoCoil gels undergo rapid softening upon 405 nm light exposure, indicated by a color change in the material, followed by full degradation of the network. In this work, we utilized the recently evolved PhoCl2c variant to provide the most complete PhoCl cleavage possible, which we found essential to forming a truly photodegradable network (*34*).

Connecting the two coil domains to the central PhoCl protein is XTEN (Block B), a pseudo-repeat protein sequence containing only A, S, T, P, G, E amino acids. XTEN was originally evolved as a protein mimic of PEG to “eXTENd” the circulating half-life of drugs in the body (*35*, *36*). As such, it was specifically designed to be high expressing and unstructured, with the restricted amino acid makeup chosen to minimize immunogenicity. In PhoCoil, we utilize a 144 amino acid variant of XTEN, split into two 72 amino acid segments by the PhoCl protein. XTEN provides flexibility to the PhoCoil protein, minimizing steric restrictions to further encourage intermolecular interactions. In this way, XTEN functions as an alternative to the widely utilized elastin-like protein (ELP) linkers seen in many recombinant protein hydrogel designs used for 3D cell culture and delivery (*6*, *18*, *26–30*, *37–41*).

Expression and purification of the PhoCoil protein yielded a solution that fluoresced green and contained protein of the expected molecular weight by mass spectroscopy (**Supplementary Figure 1**). PhoCoil also displayed the expected photocleavage activity in solution via mass spectroscopy and sodium dodecyl-sulfate polyacrylamide gel electrophoretic (SDS-PAGE) analysis, with approximately 75% of the total protein cleaving in response to visible light (**Supplementary Figures 1-2**). We hypothesize that the remaining 25% of intact PhoCoil observed in SDS-PAGE lacks a properly matured chromophore, inhibiting its ability to undergo the expected photocleavage. This hypothesis is supported by the disappearance of the mature, intact PhoCoil peak after 405 nm light exposure and concomitant appearance of an immature, intact PhoCoil peak indicated by a comparative +18 Da shift (*10*) (**Supplementary Figure 1**).

### Characterization of stiffness, shear thinning, and self-healing behavior

Mechanical characterization of PhoCoil gels was conducted via oscillatory shear rheology. At constant low strain (5%) and frequency (10 rad s^-1^) within the linear viscoelastic range, we found that gel stiffness as measured by the storage modulus could easily be modulated between 1-4 kPa by varying the weight percent of the gel (7.5 to 12.5 wt%) (**Figure 2A**). This allowed us to capture the mid-range of soft tissue stiffness in the body, which vary from 0.5-30 kPa (*42*). Frequency sweeps indicated no crossover withing the tested range of 0.1-50 rad s^-1^ (**Figure 2B**). Shear and strain sweeps demonstrated the shear thinning behavior of the gel, with the viscosity decreasing by two orders of magnitude or more as the shear rate increased from 0.1 to 50 s^-1^ (**Figure 2C**). Strain sweeps indicated a gel-sol transition, signified by a cross over in G’ and G”, for all gel weight percentages. The 7.5 wt% and 10 wt% gel-sol transitions occur around 50% strain, while the 12.5 wt% gel transition occurred at 150% strain (**Figure 2D-E**). These viscoelastic properties are typical of coiled-coil-based protein hydrogels (*43*).

**Fig. 2.**
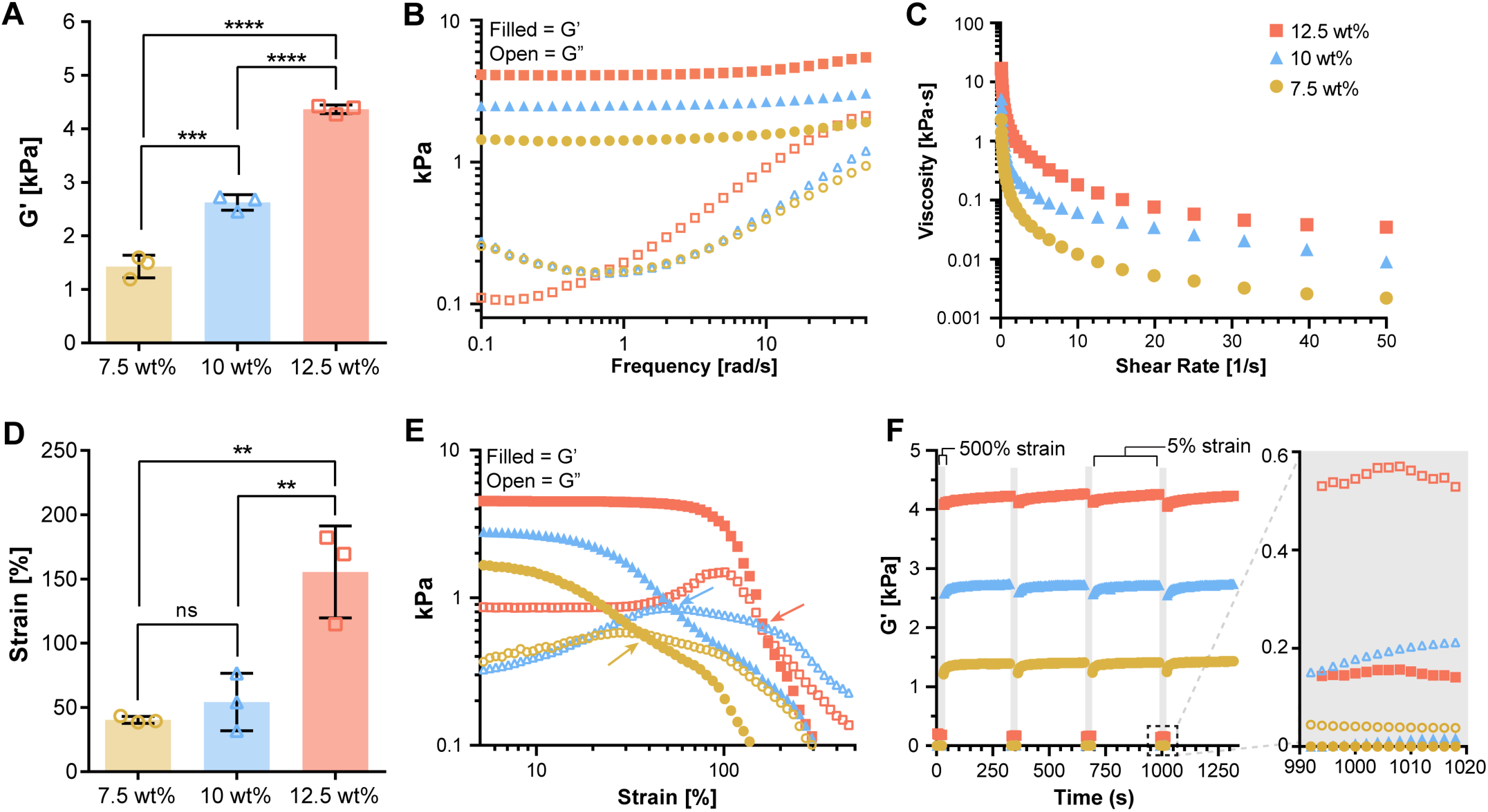
Viscoelastic properties of PhoCoil gels. (A) G’ determined from oscillatory rheology time sweeps at 25 °C, 5% strain, 10 rad s^-1^ with varied gel weight percentage. (B) A representative frequency sweep test at 25 °C with fixed 5% strain. (C) A representative viscosity profile during shear-sweep test at 25 °C. (D-E) Strain-sweep test at 25 °C with fixed 10 rad s^-1^ frequency. Panel D shows the strain crossover value indicated by the arrows in each selected representative strain sweep shown in Panel E. (F) Cyclic strain sweep test at 25 °C. Frequency was fixed at 10 rad s^-1^, strain was held at 500% in grey regions (30s) and 5% in white regions (600s). A magnified view of one high strain interval is shown to demonstrate the strain induced gel-sol transition. For all graphs, filled symbols represent G’, open symbols represent G”. All bars represent the mean ± SD of three independently formed gels for each weight percentage. Significance testing was performed using Tukey’s multiple comparisons test.

Cyclic strain sweeps were conducted to investigate the ability of PhoCoil gels to repeatedly shear-thin and self-heal via the self-associating coil domains. When the strain was increased from 5 to 500%, well above the strain crossover for all gels, G” dominated, indicating that the gels were occupying a liquid-like state (**Figure 2F**). This transition happened in under 2s, the minimum sampling interval achievable for this test. When the material was returned to 5% strain, far below the strain crossover, it quickly return to a gel state, where G’ dominates. Upon this re-gelation, stiffnesses similar to the initial G’ were achieved. All gel weight percentages were able to undergo repeated, rapid shear-thinning and complete self-healing over 4 high strain periods. These shear-thinning and self-healing behaviors indicate that PhoCoil should be easily injectable post-gelation (*2*, *44*). Injectability was confirmed by visualization of injection through a 25-gauge needle, where a stable gel is formed after exit from the needle (**Supplementary Figure 3**, **Supplementary Videos 1-2**).

### Light-induced degradation and controlled softening

We next aimed to characterize PhoCoil’s ability to soften and degrade in response to 405 nm light. To do so, we utilized a photorheometer setup that permits controlled light delivery to the gel while simultaneously measuring its mechanical properties. We found that PhoCoil gels at all weight percentages degrade in a similar “exponential decay” fashion, with first-order rate constants of 0.0293 ± 0.0009 min^-1^ (7.5 wt%), 0.024 ± 0.002 min^-1^ (10 wt%), and 0.023 ± 0.002 min^-1^ (12.5 wt%), corresponding to half-lives of 23.7 ± 0.7 min, 29 ± 2 min, and 31 ± 3 min, respectively. All weight percentages showed a plateau stiffness that was consistently 20-30% of the initial stiffness (**Figure 3A**). This initial rapid decrease in stiffness is believed to be directly related to PhoCl cleavage, which immediately weakens the gel network. The plateau stiffness that remains is hypothesized to be due to both incomplete cleavage of PhoCl, characterized in **Supplementary Figure 2** to reach a maximum of 75% cleavage, as well as entanglement of the protein chains. This rheological method only allows for the accurate capture of initial softening rather than the full gel-sol transition that would be indicated by a G’/G” crossover, as complete degradation relies on diffusion which is highly restricted in parallel-plate rheology.

**Fig. 3.**
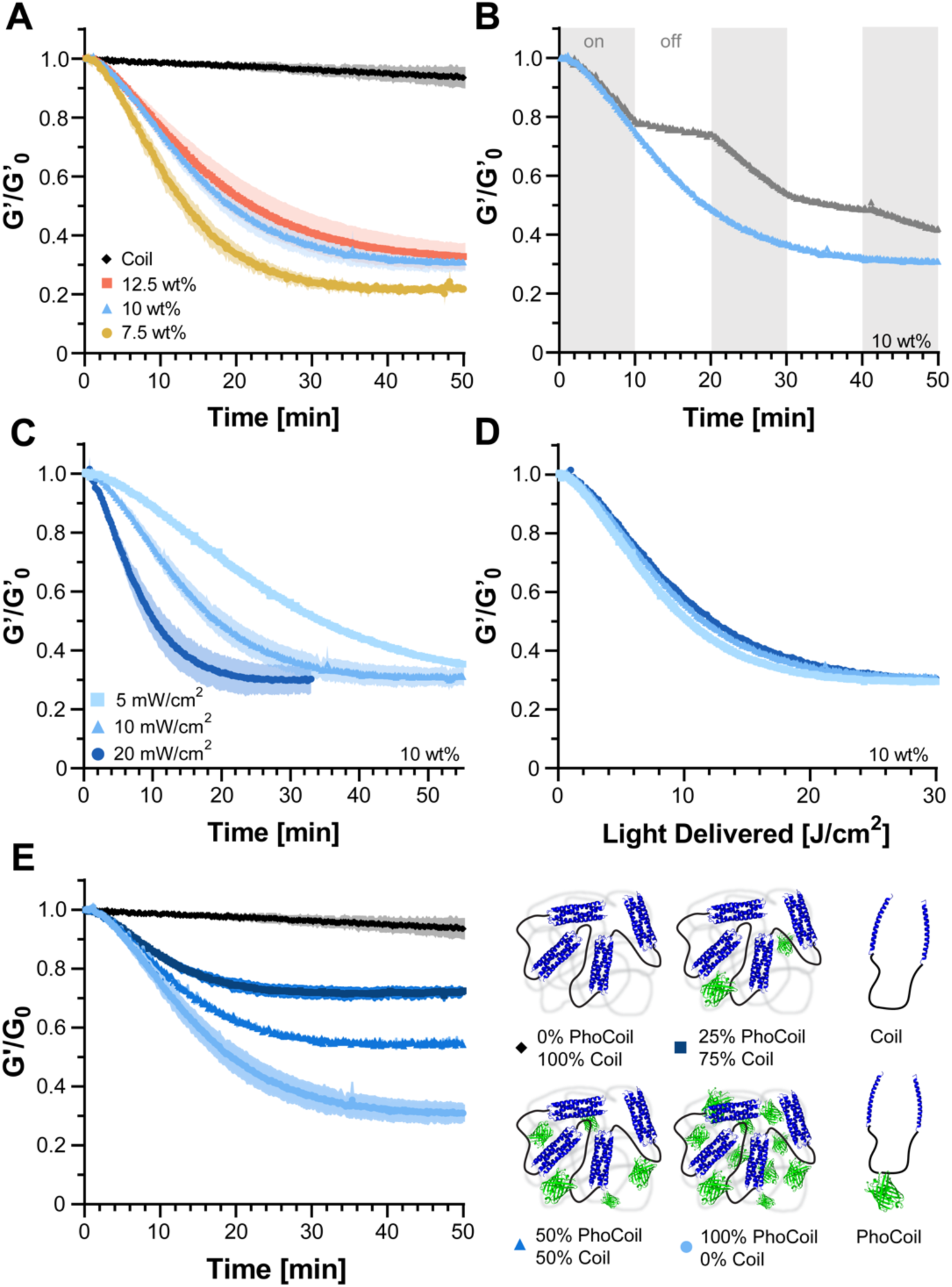
Kinetics of PhoCoil gel photodegradation. (A) Gel softening in response to 405 nm light via oscillatory rheology time sweep. Rheology was conducted at 5% strain and 10 rad s^-1^ with light intensity set to 10 mW cm^-2^ at the gel. G’_0_ was defined as the storage modulus immediately before light exposure. Three independent gels were measured for each weight percentage. Data is presented as the mean (line) ± SD (shaded area). (B) Control of endpoint stiffness by modulation of light exposure time. 10 wt% gels were exposed to constant 10 mW cm^-2^, 405 nm light (blue) or periods of 10 min light on, 10 min light off (gray). (C) Control of softening rate by modulation of light intensity. Three independent 10 wt% gels were measured for each intensity of 405 nm light. Data is presented as the mean (line) ± SD (shaded area). (D) When normalized for light dosage, experimental data from Panel C collapses onto a single curve, indicating PhoCoil photocleavage follows the same light dose-dependency for each intensity. (E) Control of endpoint stiffness by coformulation of PhoCoil with a non-light-responsive network-forming Coil protein. Three independent gels for each formulation were exposed to 10 mW cm^-2^, 405 nm light, with the total molar concentration of protein kept constant across formulations. Data is presented as the mean (line) ± SD (shaded area).

For comparison, we also tested the Coil protein developed in our previous work (*9*), which has an identical structure to PhoCoil but lacks the PhoCl protein. As expected, Coil shows minimal change in stiffness when exposed to 405 nm over the same period of time, indicating that the photoresponsivity of PhoCoil stems directly from its mid-block PhoCl protein. The minimal softening of Coil captured over the course of photorheology is also seen without light exposure over the same interval and is expected to be a result of the gel swelling and eroding at the edges, as measurements are taken with the gel submerged in phosphate-buffered saline (PBS) (**Supplementary Figure 4**).

To further probe PhoCoil’s light responsive properties, we exposed 10 wt% PhoCoil gels to intermittent and varying intensities of light. Here we showed that PhoCoil softening is directly controlled by the time and intensity of light exposure, and can be modulated as such. PhoCoil softening can be stopped before completion by removing light, resulting in immediate plateauing of the gel stiffness (**Figure 3B**). The rate of gel softening can also be modulated by changing the intensity of the light, with higher intensities leading to more rapid softening to the same plateau value (**Figure 3C**). Notably, the extent of softening scales directly with the total light dosage delivered (given in J cm^-2^) regardless of the intensity of the source, with an intensity adjusted rate constant of 0.0451 ± 0.0005 cm^2^ × J^-1^ and half-life of 15.4 ± 0.2 J × cm^2^ for a 10 wt% gel (**Figure 3D**). This relationship allows for simple calculation of the intensity of light and time of light exposure that can be used to reach a desired final stiffness if the initial gel stiffness is known.

While we have demonstrated that partial softening can be achieved in a dose-dependent manner, there are some scenarios in which it may be useful to fix the minimum final stiffness that the material can be softened to. We hypothesized that this could be achieved by co-formulating PhoCoil with our non-light-responsive Coil protein. These co-formulated gels soften to intermediate final stiffnesses, with higher percentages of Coil leading to less softening despite receiving the same dosage of light (**Figure 3E**).

To capture the complete degradation of PhoCoil gels, we turned to bulk degradation studies. PhoCoil gels at 7.5, 10, and 12.5 wt% were formed in microcentrifuge tubes and covered with excess PBS. The gels are initially green in color due to the inherent green fluorescence of PhoCl. Upon exposure to 405 nm light, the gels undergo a visible color change to orange as a result of the cleavage and rearrangement of PhoCl’s chromophore, which occupies a red state prior to separation of the two halves of the protein (*10*) (**Figure 4A**). We hypothesize that due to the molecular crowding within the hydrogel this separation is slowed, allowing the red state to persist longer than in solution. The gel then primarily erodes from the surface, where cleaved protein can more easily escape the network. Complete degradation is achieved in response to 405 nm light within 4 to 20 hours under these conditions, with increasing doses of light decreasing the time to complete degradation (**Figure 4A-B**). For gels kept in ambient light only (0 min light), no noticeable color change occurred through the duration of this experiment, indicating that PhoCl is resistant to photocleavage in ambient light. However, PhoCoil is susceptible to surface erosion in the absence of directed photocleavage, as the network is only bound by dynamic, physical interactions. As such, coil exchange at the surface of the gel makes it possible for uncleaved polypeptides to diffuse away from the network while their coils are in an unbound state, as demonstrated with the Coil networks in previous work (*9*). PhoCoil gels that are not exposed to 405 nm light persist for 3 days or longer under these conditions (**Figure 4A**, **Supplementary Figure 5**). Direct 405 nm light exposure significantly increased the rate of degradation; with 90 minutes of exposure to visible light (405 nm, 10 mW cm^-2^), PhoCoil gels have half-lives that are 20-30 times shorter than those that are kept in ambient light (**Figure 4B, Supplementary Table 1**). As expected, increasing the weight percentage of the network increases the average time to complete degradation for all light conditions.

**Fig. 4.**
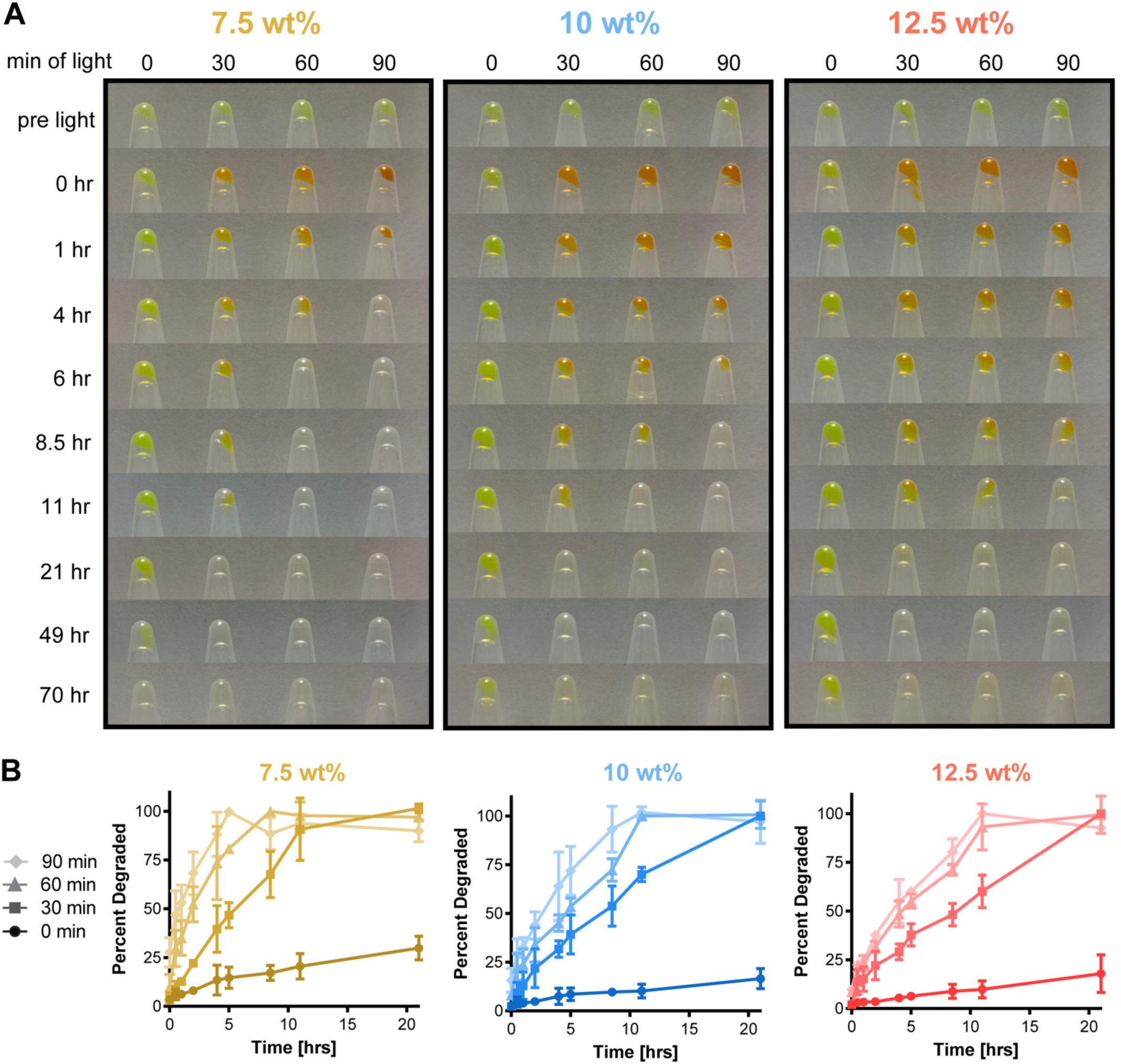
Bulk degradation of PhoCoil gels. (A) Degradation of varying gel weight percentages in response to increasing time of light exposure. 25 μL gels covered with PBS were exposed to 10 mW cm^-2^, 405 nm light for 0, 30, 60, or 90 min. Images of remaining gel in inverted tubes were obtained at the indicated intervals post-exposure. (B) Quantification of gel degradation rates. Samples of PBS above each gel were analyzed by BCA assay to determine protein content as a measure of the percentage of gel degraded at each interval post exposure. Three independent gels were tested for each combination of weight percentage and light exposure time. Data shown are mean ± SD.

### Spatiotemporal control of gel softening and degradation

One significant advantage of light as a stimulus is the ease of controlling its delivery spatiotemporally (*11*). Encouraged by our successful demonstration of PhoCoil’s ability to photodegrade, we sought to control material dissolution in user-defined patterns (**Figure 5A)**. Here, we use photomask-based lithography to spatially control light exposure, resulting in clear patterns with micron-scale resolution (**Figure 5B-C**). The resulting patterns can easily be visualized with confocal microscopy by using the intrinsic color switch of PhoCl after cleavage. Intact, non-light exposed areas have strong green fluorescence, while the photocleaved areas have increased red fluorescence.

**Fig. 5.**
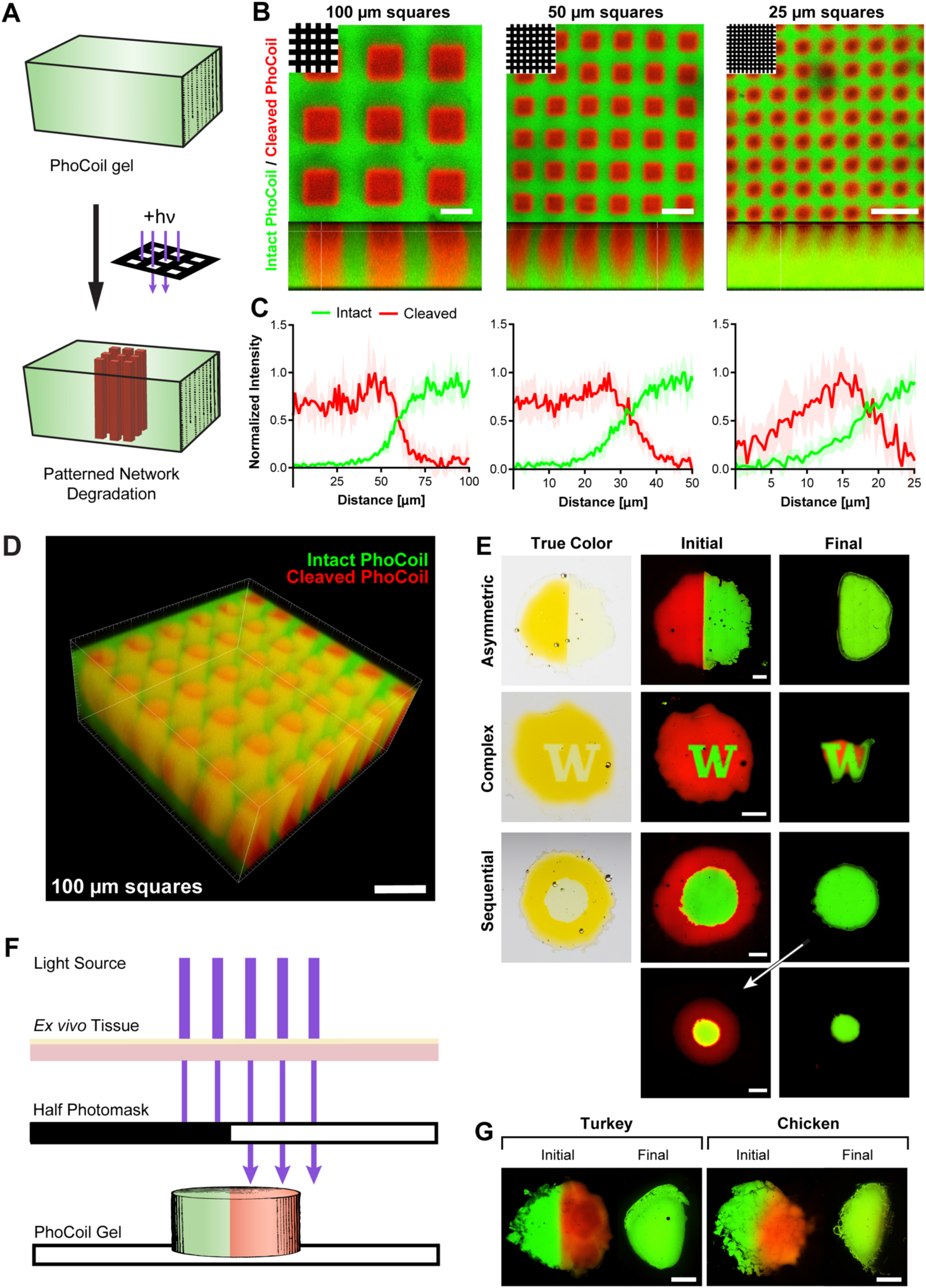
Photopatterning and spatially controlled degradation of PhoCoil gels. (A) PhoCoil gels can be patterned using photomasks to restrict 405 nm light to desired areas. (B) Photomask-based patterning of grids of varying sizes onto PhoCoil gels. Gels were exposed to 60 min of 10 mW cm^-2^, 405 nm light through a grid photomask with light passing through 100, 50, or 25 μm-wide squares (top left). Images were obtained in the xy (top) and xz (bottom) planes via confocal microscopy. Scale bars are 100 μm. (C) Intensity in red and green channels was quantified via ImageJ to demonstrate changes in resolution of patterning. Intensity was quantified from the center of a red square to the center of a green line at 5 random locations. Data shown is mean ± SD. (D) 3D reconstruction of the 100 μm grid patten throughout the thickness of a gel. Scale bar is 250 μm. (E) Spatially controlled degradation of photomask patterned gels. Gels were exposed to 60 min of 10 mW cm^-2^, 405 nm light through a photomask, then photographed (true color) or confocal imaged (initial). Gel degradation was allowed to occur in excess PBS, and gels were confocal imaged to visualize degradation (final). Scale bars are 1 mm. (F&G) Light-based degradation of PhoCoil gels through *ex vivo* tissue. The left half of each gel was covered with a photomask, and the entire gel was placed under 1 mm-thick deli turkey or chicken skin. Gels were exposed to 5 min of high-intensity 405 nm light through the tissue mimic, confocal imaged to visualize the pattern (initial), and left to degrade in excess PBS prior to final fluorescent imaging. Scale bars are 1 mm.

Photomask-based patterning can be achieved throughout a 500 µm thick gel with good resolution at the ∼100 µm scale (**Figure 5D**). However, as pattern details get smaller, scattering of light within the hydrogel results in a reduction in pattern crispness and depth (**Figure 5B-C**).

We next used photomask-based lithography to spatiotemporally control complete gel degradation. To do so, we patterned gels with a variety of photomasks, then submerged them in a PBS bath with intermittent confocal imaging to capture various stages of degradation. Imaging showed that the light-exposed areas of the gels quickly degraded, while the unexposed areas persisted (**Figure 5E**). Patterned degradation could be achieved in several configurations, including a half gel, central “W”, and central circle. Sequential light exposures permitted staged degradation of different sections of the material.

To determine if controlled photodegradation could be achieved in tissue, we utilized *ex vivo* tissue (deli turkey or skin-on chicken) as a barrier for light to pass through to reach the PhoCoil gel (**Figure 5F**). Degradation of the unmasked sections of these gels was easily achieved under thin tissue sections (∼1-2 mm), demonstrating that PhoCoil gels may be useful for controlled therapeutic delivery at low injection depths (**Figure 5G**, **Supplementary Figure 6**).

### Cell viability through encapsulation, injection, and light-mediated release

Given our success in demonstrating both injectability and photodegradation of PhoCoil, we lastly aimed to assess its potential utility in cell delivery. In our previous work with the Coil system, we demonstrated that Coil gels support high cell viability through encapsulation and injection, both *in vitro* and *in vivo* (*9*). As the PhoCoil gels are formed in an identical manner to Coil gels, it remains straightforward to encapsulate cells in PhoCoil gels. Cells in suspension (viability >99%) are combined with the PhoCoil protein to yield encapsulated cells, which retained 85% viability at 1.5 hours post-encapsulation (**Figure 6A-D**). For cells encapsulated in PhoCoil and subsequently injected through a 25-gauge needle, 79% viability was observed (**Figure 6B,E**).

**Fig. 6.**
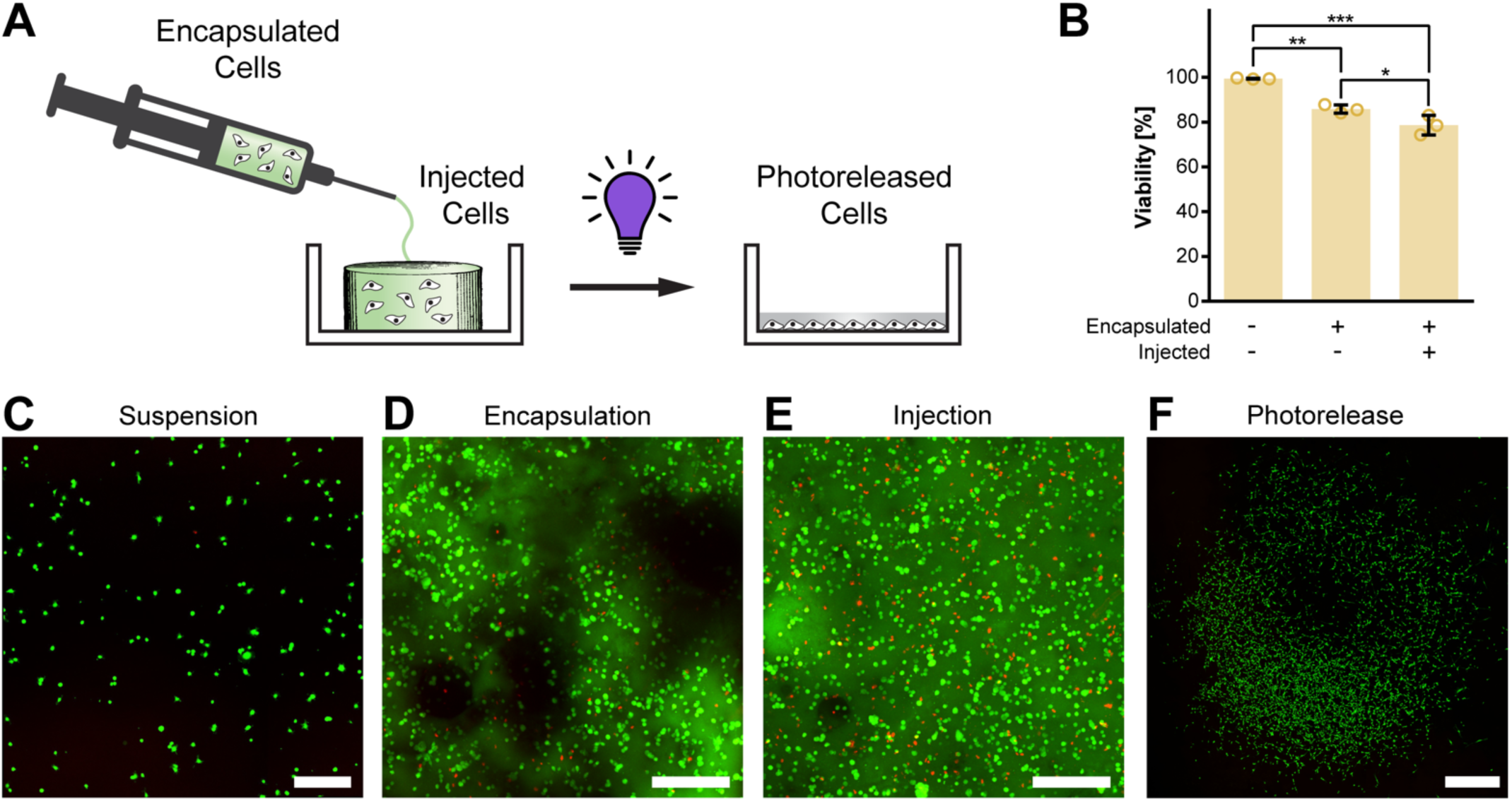
Viability of PhoCoil-encapsulated fibroblasts. (A) 10T1/2 fibroblasts are encapsulated in gel, injected into a well plate, and released from the gel via 405 nm light. (B) Viability of 10T1/2 fibroblasts throughout mock processing (in suspension, after encapsulation in gel, and after injection in gel) for therapeutic cell delivery in PhoCoil gels. All bars represent the mean ± SD of three independent gels for each condition. Significance testing was done using Tukey’s multiple comparisons test. (C) Representative confocal microscopy image of Live/Dead analysis of 10T1/2 fibroblasts in suspension. Scale bar is 250 μm. (D) Representative confocal microscopy image of Live/Dead analysis of 10T1/2 fibroblasts post-encapsulation in a 7.5 wt% PhoCoil gel. Image is a maximum projection of a z-stack through a 0.5 mm-thick gel. Scale bar is 250 μm. (E) Representative confocal microscopy image of Live/Dead analysis of 10T1/2 fibroblasts post-injection in a 7.5 wt% PhoCoil gel. Image is a maximum projection of a z-stack through a 0.5 mm thick gel. Scale bar is 250 μm. (F) Representative confocal microscopy image of Live/Dead-stained 10T1/2 fibroblasts released from a PhoCoil gel. Cells were subjected to encapsulation and injection, followed by release from the gel into a cell culture dish using 405 nm light exposure. Two days following release, cells were stained and imaged to assess their ability to survive and proliferate after such handling. Scale bar is 1 mm.

Unlike our previously developed Coil system, cells encapsulated in PhoCoil hydrogels can be subsequently released using visible light to liberate single-cell suspensions at a user-defined time point. To demonstrate this, encapsulated cells were exposed to 405nm light to induce gel photodegradation. Photoreleased cells were then left for 2 days to encourage readherence to the dish and proliferation prior to imaging. These cells retained their proliferative capacity and expanded to cover a tissue culture plate over this 2-day period (**Figure 6F**). Given PhoCoil’s ability to support high cell viability throughout injection and photodegradaton, it is poised for applications in therapeutic cell delivery, where it can provide a protective matrix for cells that will degrade slowly overtime or quickly after light exposure.

## Discussion

In this work, we have showcased the unique properties of PhoCoil, a single-component recombinant protein hydrogel system that is biocompatible, injectable, and photodegradable. The molecular design of the PhoCoil protein allowed us to preemptively engineer in our desired properties to realize this dynamic material. PhoCoil forms supramolecular hydrogels that with tunable stiffnesses spanning that of many native tissues, and its injectability primes it for applications in minimally invasive, controlled therapeutic delivery. Furthermore, the photoresponsivity of PhoCoil allows for its softening and degradation to be bioorthogonally and spatiotemporally controlled in a precise and dose-dependent manner. We showed that this photodegradation can be achieved under thin layers of *ex vivo* tissue, suggesting its potential utility for transdermally triggered dissolution. Furthermore, we demonstrated that PhoCoil gels are cytocompatible, opening up their application to cell-based therapeutics and regenerative medicine applications. Collectively, this work provides proof-of-concept evidence of PhoCoil’s utility in therapeutic cell delivery.

To take advantage of PhoCoil’s photodegradability for rapid material dissolution, it is best positioned for intradermal or subcutaneous therapeutic delivery, or for applications in more optically transparent tissues (e.g., eyes). However, significant advances in protein engineering and *de novo* protein design may enable the development of further red-shifted variants of PhoCl (*45*, *46*). This would be particularly useful, as longer wavelengths of light penetrate deeper through tissue (*47*, *48*). Furthermore, continued development of tools that allow shorter wavelengths of light to penetrate deeper into the body, such as upconverting nanoparticles, implantable LEDs, and optical waveguides (*48–51*), offer exciting opportunities for combinatorial approaches that would further enable *in vivo* regulation of PhoCoil and other light-responsive materials.

As cell-based therapies continue to be a focus of next-generation therapeutic approaches, hydrogels will surely play a role in their development and effective implementation (*52*). Novel, recombinant protein-based materials, such as PhoCoil, are well positioned to be utilized for translational therapies given their perfectly defined, tunable, and bioresorbable nature. Recent advances in *de novo* protein design have also enabled development of self-associating proteins not seen in nature, opening up new possibilities in the design of genetically encoded protein biopolymer networks (*53*). The recent emphasis in biotech on protein-based therapeutics promises to develop more efficient and cost-effective ways to screen protein expression conditions and achieve large scale protein expression, which will further enable the usage of such recombinant protein materials (*54*, *55*).

## Materials and Methods

### Cloning

A gBlock (IDT) containing the sequence encoding 6xHis-RGD-Coil-XTEN-PhoCl2c-XTEN-Coil-RGD-6xHis, herein referred to as PhoCoil, was inserted into the pET21 vector via Gibson assembly (NEB). This plasmid is available through Addgene (ID 220444). The resulting mixture was transformed into TOP10 *E. coli* and the sequence confirmed by Sanger sequencing (Genewiz). The plasmid was then transformed into electrocompenent BL21(DE3) *E. coli* for protein expression work.

### Protein Expression

BL21(DE3) *E. coli* containing the pET21 PhoCoil plasmid was grown overnight in Lysogeny Broth (LB). Fresh LB medium was combined with carbenicillin and inoculated 1:200 with the overnight culture. The culture was grown at 37°C and 210 rpm and the OD600 monitored until 0.4-0.6, at which point 0.5 mM isopropyl β-D-1-thiogalactopyranoside (IPTG) was added and the culture was moved to 18°C, 210 rpm for 24 hrs in the dark. Cells were harvested by centrifugation of 500 mL of culture at 4000 xg for 20 min at 4°C. Cell pellets were frozen at −80°C at least overnight.

### Protein Purification

Cell pellets, each derived from 500 mL of culture, were resuspended in 40 mL of lysis buffer (50 mM NaCl, 20 mM Tris, 10 mM imidazole, pH 8) supplemented with 1mM phenylmethylsulfonyl fluoride (PMSF). Resuspended cells were placed in an ice water bath and sonicated at 30% amplitude for 6 min of total on time in 1s on, 2s off intervals (Fisherbrand, Model 505 Sonic Dismembranator, 0.5-inch probe). Cell lysate was centrifuged at 10,000 xg for 20 min at 4°C, and the supernatant was collected and stored at 4°C for 7 days to encourage full chromophore maturation of PhoCl. The clarified lysate was again centrifuged at 10,000 xg for 20 min at 4°C and the supernatant collected for purification. Ni-NTA HisTag affinity chromatography was conducted using an ÄKTA Pure 25L with a 5-mL HisTrap column (Cytiva). 40 mL of clarified cell lysate was loaded onto the column. The column was then thoroughly washed (50 mM NaCl, 20 mM Tris, 15 mM imidazole, pH 8) and the protein eluted (100 mM NaCl, 40 mM Tris, 500 mM imidazole, pH 7.5). Collected elution fractions containing protein were desalted into water using a 50 mL 26/10 desalting column (Cytiva) on the ÄKTA. The column was equilibrated with deionized water, and 13 mL of the collected elution was loaded, followed by elution with deionized water. Only the protein peak (high A280, low conductivity) was collected, flash frozen in liquid nitrogen, and lyophilized for 3 days. Lyophilized protein was stored in parafilm-wrapped tubes at −20°C.

### Gel Formation

Gels were formed by resuspending lyophilized protein in sterile 10X PBS, pH 7.4 at 7.5, 10, or 12.5 wt%. Each mg of protein was assumed to take up 1 μL of volume in the mixture. Immediately after PBS addition, tubes were placed at 37°C for 5 min. The mixture was then thoroughly stirred with a pipette tip to break up chunks of protein and returned to 37°C for 20 min, at which point a cohesive gel was observed. The gels were centrifuged at 10,000 xg for 5 min to remove trapped bubbles and placed at 4°C on a rocker overnight prior to use.

### Rheology

Rheology was performed on an Anton Paar Physica MCR 301 using an 8 mm-diameter parallel-plate geometry with a 0.5 mm gap, at 25°C. Approximately 60 μL of preformed gel was loaded and covered with mineral oil to prevent evaporation. The following testing routine was performed on each gel: Time sweep (10 rad s^-1^, 5% strain, 30 min), Frequency sweep (0.1-50 rad s^-1^, 5% strain), Time sweep (10 rad s^-1^, 5% strain, 10 min), Strain sweep (10 rad s^-1^, 0.1-500% strain), Time sweep (10 rad s^-1^, 5% strain, 10 min), Cyclic strain sweep (10 rad s^-1^, 5% strain for 5 min then 500% strain for 30s, repeated 4 times total), Time sweep (10 rad s^-1^, 5% strain, 10 min), Shear sweep (0.1-50 s^-1^ shear rate). G’ values were determined by averaging all measurements in the initial time sweep. Crossover strain values were determined through linear interpolation.

### Photorheology

Photorheology was performed on an Anton Paar Physica MCR 301 using an 8 mm-diameter parallel-plate geometry and a custom, optically transparent lower plate with a fitting for a light guide. All measurements were performed with a 0.5 mm gap at ambient temperature, with frequency held at 10 rad s^-1^ and strain at 5%. Preformed gels were covered with excess PBS pH 7.4 and allowed to swell for 1-2 hrs. Gels were then loaded onto the rheometer and trimmed with plastic tweezers to the edge of the geometry. The well in the custom plate was filled with 5 mL of PBS to prevent the gel from drying out or heating up significantly during the run. The gel was allowed to equilibrate and stiffness plateau for 15 min, at which point the light source (Mightex WheeLED, 405nm, 5-20 mW cm^-2^ at gel) was turned on for 50 min to capture gel softening. For the interval light exposure, the light was turned on for 10 min, then off for 10 min for a total of 50 min. The final G’ value prior to light exposure was set as G’_0_.

### Bulk Degradation Study

Gel degradation behavior was assessed in PBS with and without light exposure. Preformed 25 μL gels in 1.5 mL microfuge tubes were washed with 1 mL of PBS twice, then covered with 1 mL of PBS, pH 7.4. Gels were then exposed to 405 nm light at 10 mW cm^-2^ for 0, 30, 60, or 90 min, with tubes rotated 180° every 15 min to ensure uniform light exposure. Tubes were then placed in a shaker incubator at 37°C and 150 rpm, with the tubes oriented parallel to the shaker plate. Images of the gels and aliquots of the PBS above the gels were taken over the course of 3 days. Each time, fresh PBS was added to each tube to maintain a constant volume. The protein concentration of each sample was determined by BCA assay (Thermo Scientific, Pierce BCA Protein Assay Kit) relative to a bovine serum albumin (BSA) standard curve. Data was adjusted to account for evaporation over the course of the study based on the volume of liquid remaining at the end of the study and assuming linear volume loss. Values were normalized to from 0-100%, with the 100% point selected based on cross referencing the data with the provided images of the gels. Half-lives were determined from the first order rate constant, calculated for each replicate, and averaged.

### Photomask-based Patterning and Degradation

Preformed gels were divided into 5-10 μL volumes using plastic tweezers and placed onto a glass slide. Rubber spacers (0.5 mm thick) were placed on each side of the gel and a coverslip was placed on top to flatten the gel into a disc. The remaining empty space between the slide and coverslip surrounding the gel was filled with PBS, pH 7.4 to keep the gel hydrated. The mounted gel was oriented with the coverslip touching the photomask and the glass slide covered with a piece of black rubber in order to prevent residual light passing through the gel from backscattering. A 405 nm light source was oriented to shine through the photomask and onto the gel with a power of 10 mW cm^-2^ at the gel surface. Light exposure was maintained for 1 hr, at which point the gels were imaged on a Leica Stellaris 5 confocal microscope to visualize the resultant pattern. Intact PhoCoil (green) was visualized with an excitation wavelength of 489 nm and an emission detection range of 494-572 nm. Cleaved PhoCoil (red) was visualized with an excitation wavelength of 587 nm and an emission detection range of 592-750 nm. To quantify the patterning resolution of the grids, images were analyzed in ImageJ. A line was drawn from the center of light exposed region to the center of a non-light exposed region, and the value of the green and red channels over this line were determined using the plot profile tool. This was repeated 5 times per grid size at random locations in the image. For 3D images, z-stacks were reconstructed in ImarisViewer. For photodegradation, patterned gels were imaged immediately after light exposure, then the entire slide was placed in a bath of PBS in an incubator shaker at 37°C and 100 rpm. Gels were intermittently imaged to capture degradation of the light exposed areas of the gel.

### Ex Vivo Photodegradation

Preformed gels were prepared between glass slides as detailed above. Half of the gel was covered with a rubber photomask, and the whole gel was covered with the indicated tissue (deli turkey or skin-on chicken). A 405 nm light source was setup approximately 1mm above the tissue. The following condition were used: 1mm of deli turkey, 150 mW at source, 5 min exposure; 2mm of deli turkey, 150 mW at source, 10 min exposure; 2mm of deli turkey, 25 mW at source, 60 min exposure; chicken skin alone (1 mm), 125 mW at source, 5 min exposure; 2mm of chicken with skin on 125 mW, 15 min exposure. The tissue was intermittently sprayed with deionized water to keep it from drying. Immediately after light exposure, gels were imaged by confocal microscopy as noted above. Each slide was then placed in a bath of PBS overnight at ambient temperature, and confocal image of the remaining gel was captured.

### Cell Viability

Cell viability during encapsulation and injection was assessed using 10 T1/2 fibroblasts. Cells (passage 8) were cultured in dishes until ∼70% confluent, then treated with TrypLE to obtain a single cell suspension. Cells were pelleted, then suspended in FluoroBrite medium supplemented with 10% FBS, 1x penicillin-streptomycin, and 25 mM HEPES at >2 million cells mL^-1^. For the suspension condition (- encapsulation, -injection), cell suspension was dispensed into a tissue culture dish and stained with 2 mM Calcein AM and 1 mM Ethidium Homodimer for 30 minutes prior to imaging. For the gel conditions (+encapsulation, ±injection), cell suspension and fresh media were used to resuspend lyophilized PhoCoil to obtain 7.5 wt% gels with 2 million cells mL^-1^ of gel. After combining, the mixture was placed at 37 °C for 10 min, then mixed by stirring with a pipette tip to ensure homogenous suspension of cells in the gel. This was repeated 3 more times, then the mixture was briefly centrifuged to gather the gel at the bottom of the tube. Gels were dispensed by positive displacement pipette (-injection) or through a Hamilton syringe with a 0.4”-long, 20-gauge needle (+injection) into rubber molds placed in 30 mm dishes. These molds had circular cavities 0.5 mm deep and 10 μL in volume to hold the gels. Three gels (10 μL) were placed in each dish and covered with 8 mM Calcein AM and 2 mM Ethidium Homodimer in PBS for live/dead analysis. Gels were incubated at 37°C in stain for 1.5 hours before imaging on a Leica Stellaris Confocal Microscope to visualize cells. Z-stacks thorough the thickness of the center of each gel were obtained, and maximum intensity z-projections of each were used to quantify cell viability by manually counting live (green) and dead (red) cells. For light-mediated cell release, molds were removed from around the gels which were then covered with 100 μL of fresh media. 405 nm light (100 mW at source) was positioned directly above the dish lid over the gel for 10 min. Photodegraded gels were then covered with additional Dulbecco’s Modified Eagle Medium (DMEM) and incubated for 2 days to allow for cell growth. After 2 days, the media was supplemented with 2 mM Calcein AM and 1 mM Ethidium Homodimer and cells were allowed to stain for 30 minutes prior to imaging.

### Statistical Analysis

In all figures, n=3 replicates were used, where each replicate was an independent gel. Where error bars are shown, all values indicate the mean ± SD. Significance was determined using Tukey’s multiple comparison test in GraphPad Prism.

## Acknowledgements

The authors would like to acknowledge Dale Whittington for training and maintenance of the UW Mass Spectrometry Center. Authors also acknowledge helpful advice, feedback, and training from Teresa Rapp, Irina Kopyeva, Ross Bretherton, Mary O’Kelly Boit, Jenny Bennett, and Ryan Francis.

## Funding

National Science Foundation Graduate Research Fellowship Program DGE-2140004 (NEG) NIH Maximizing Investigators’ Research Award R35GM138036 (CAD)

Part of this work was conducted with instrumentation provided by the Joint Center for Deployment and Research in Earth Abundant Materials (JCDREAM)

## Author Contributions

Conceptualization: CAD, NEG

Methodology: NEG

Investigation and Analysis: NEG

Resources: CAD, NEG

Visualization: NEG

Supervision: CAD

Writing – original draft: CAD, NEG

Writing – review and editing: CAD, NEG

## Data and Materials Availability

All data are available in the main text or the supplementary materials.

