## Supplementary Information for "PhoCoil: An Injectable and Photodegradable Single-component Recombinant Protein Hydrogel for Localized Therapeutic Cell Delivery"

**Supplementary Materials**

**PhoCoil Amino Acid Sequence**

Coil domain  PhoCl2c  XTEN  GS Linker  RGD  6X HisTag

MRGSHHHHHHEFGRGDSVDGSGSGSGSGSGAPQMLRELQETNAALQDVRELLRQAVKEITFLK
NTVMESDASGSGSGSGSGSGSLDGSPAGSPTSTEEGTSESATPESGPGTSTEPSEGSAPGSPAGS
PTSTEEGTSTEPSEGSAPGTSTEPSEGSAKLEFVIPDYFKQSFPEGYSWERSMTYEDGGICIATNDIT
MEGDSFINKIHFQGTNFPPNGPVMQKRTVGWEASTEKMYERDGVLKGDVKMKLLLKGGGHYRGDY
RTTYKVKQKPVKLPDCHFVDHRIEILSHDKDYNKVKLYEHAVAKTSTDSMDELYKGGSGGMVSKGEE
TITSVIKPDMKNKLRMEGNVNGHAFVIEGEGSGKPFEGIQTIDLEVKEGAPLPFAYDILTTAFHYGNRV
FTKYPRGGGQLPGTSESATPESGPGSEPATSGSETPGSEPATSGSETPGSPAGSPTSTEEGTSESA
TPESGPGTSTEPSEGSAPLDGSGSGSGSGSGAPQMLRELQETNAALQDVRELLRQAVKEITFLK
NTVMESDASGSGSGSGSGSGSGRGDSHHHHHH

**PhoCoil Nucleotide Sequence (5'→ 3')**

ATGCGTGGCAGCCACCATCACCATCACCATGAGTTTGGCCGCGGAGATTCAGTAGATGGTTCGG
GCTCTGGATCAGGTTCGGGTTCGGGTAGTGGGGCCCCACAGATGCTTCGCGAGCTCCAAGAGA
CAAACGCGGCCCTGCAAGACGTACGTGAGCTTCTCCGCCAGGCAGTAAAAGAAATTACATTTCT
GAAGAATACTGTTATGGAAAGCGACGCCTCAGGTTCGGGTTCTGGGTCAGGCAGCGGCTCCGG
CTCATTGGATGGGTCGCCAGCAGGTAGTCCAACGAGTACCGAGGAAGGTACCTCAGAAAGCGC
CACTCCAGAATCAGGTCCAGGTACAAGCACTGAGCCTTCGGAAGGGTCTGCCCCCGGCTCCCC
AGCGGGCAGTCCCACGTCAACAGAGGAAGGCACGAGTACTGAGCCTTCAGAAGGGTCAGCTCC
AGGGACATCAACGGAACCTAGTGAAGGCAGCGCCAAACTGGAATTCGTGATCCCTGACTACTTC
AAGCAGAGCTTCCCCGAGGGCTACAGCTGGGAGCGCAGCATGACCTACGAGGACGGCGGCAT
CTGCATCGCCACCAACGACATCACAATGGAGGGGGACAGCTTCATCAACAAGATCCACTTCCAG
GGCACGAACTTCCCCCCCAACGGCCCCGTGATGCAGAAGAGGACCGTGGGCTGGGAGGCCAG
CACCGAGAAGATGTACGAGCGCGACGGCGTGCTGAAGGGCGACGTGAAGATGAAGCTGCTGCT
GAAGGGCGGCGGCCACTATCGCGGCGACTACCGCACCACCTACAAGGTCAAGCAGAAGCCCG
TAAAGCTGCCCGACTGCCACTTCGTGGACCACCGCATCGAGATCCTGAGCCACGACAAGGACT
ACAACAAGGTGAAGCTGTACGAGCACGCCGTGGCCAAGACTTCCACCGACAGCATGGACGAGC
TGTACAAGGGTGGCAGCGGTGGCATGGTGAGCAAGGGCGAGGAGACCATTACAAGCGTGATCA
AGCCTGACATGAAGAACAAGCTGCGCATGGAGGGCAACGTGAACGGCCACGCCTTCGTGATCG
AGGGCGAGGGCAGCGGCAAGCCCTTCGAGGGCATCCAGACGATTGATTTGGAGGTGAAGGAG
GGCGCCCCGCTGCCCTTCGCCTACGACATCCTGACCACCGCCTTCCACTACGGCAACCGCGTG
TTCACCAAGTACCCACGGGGCGGCGGCCAATTGCCGGGAACATCAGAATCTGCTACACCAGAA
TCGGGCCCTGGGTCAGAACCGGCCACTTCGGGTTCAGAAACACCTGGCAGTGAACCTGCTACT
AGTGGCTCGGAAACCCCAGGGTCCCCAGCCGGTTCCCCTACCTCGACAGAAGAGGGTACATCG
GAGTCCGCGACACCTGAATCAGGGCCAGGAACAAGCACAGAACCGTCGGAAGGGTCTGCCCCT
CTGGATGGAAGCGGTTCAGGCTCGGGTTCGGGTAGCGGTTCGGGCGCACCCCAGATGTTGCG
CGAGCTTCAAGAGACTAACGCAGCCTTACAAGACGTACGCGAGCTGCTGCGCCAGGCCGTGAA
AGAAATTACCTTCTTAAAGAATACGGTTATGGAAAGTGACGCCTCAGGGAGTGGTAGTGGTTCG
GGCAGCGGGTCAGGCTCTGGCCGCGGCGATTCACACCATCACCATCACCATTAA

*Note: PhoCoil plasmid is available at Addgene (ID 220444).*

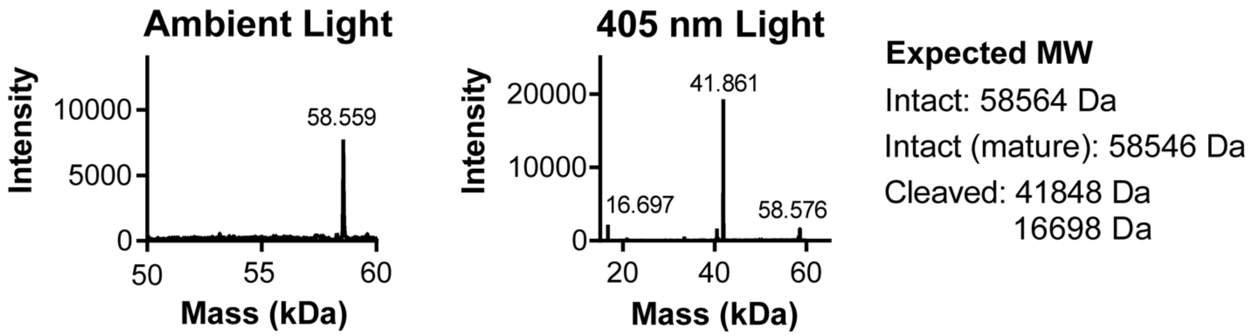


**Supplementary Fig. 1. Mass spectrometry of PhoCoil before and after 405 nm light exposure.** In ambient light conditions (left), the main peak corresponds to intact PhoCoil with a mature chromophore. The actual mass is ~13 Da higher than the mass expected from the primary amino acid sequence alone when accounting for a -18 Da shift for chromophore maturation. However, this +13 Da shift matches precedent from the coiled-coiled XTEN gels that this work was based on (*1*). After exposure to 405 nm light (10 mW $cm^{-2}$ for 60 min), the main peak corresponds to the larger fragment of cleaved PhoCoil (which retains the +13 Da mass shift from expected). The two additional peaks correspond to the smaller fragment of cleaved PhoCoil (left) and residual intact PhoCoil with an immature chromophore (right), indicated by a +18 Da shift from mature PhoCoil.

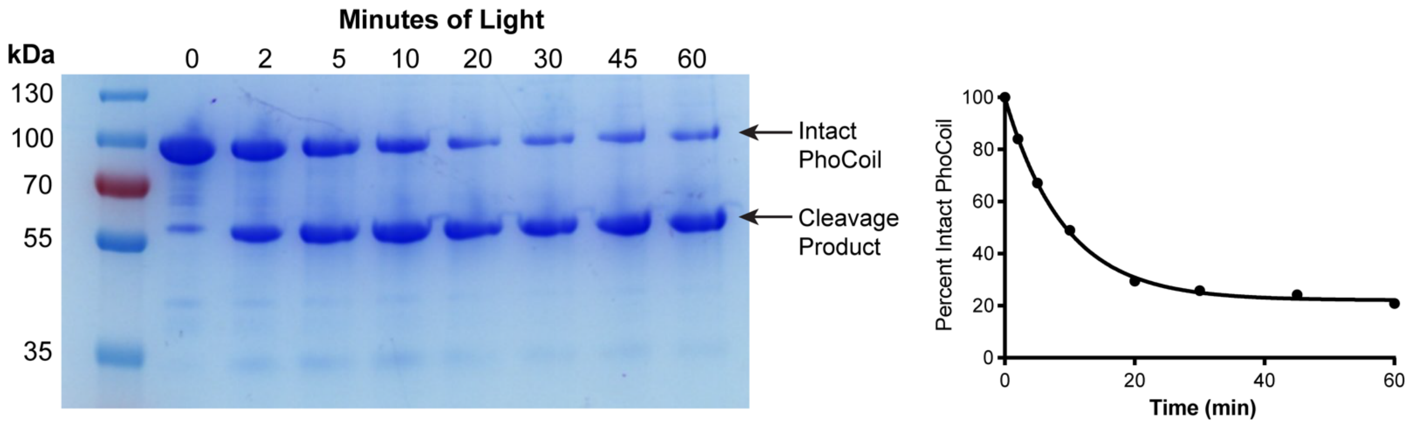


**Supplementary Fig. 2. Analysis of PhoCoil cleavage in solution.** A 1 mg $mL^{-1}$ solution of PhoCoil, too dilute to form a gel, was subjected to 10 mW $cm^{-2}$ of 405 nm light for 0-60 min. Samples were obtained at various intervals and run on SDS-PAGE with reducing and denaturing conditions. The acrylamide gel was stained with Coomassie blue to visualize protein bands. Note that it is expected that PhoCoil proteins run larger than their actual size on SDS-PAGE due to the inclusion of XTEN sequences, which lack hydrophobic amino acids and are expected to bind SDS less strongly because of this (*2*). The efficiency of PhoCoil cleavage was quantified using ImageJ to calculate the intensity of the intact PhoCoil bands in each sample. The percentage of intact PhoCoil remaining was determined by dividing each intensity by the original band intensity at 0 min, and the data was fit to an exponential decay formula. Residual intact PhoCoil is expected to be mostly in an immature state, where a lack of chromophore formation prevents cleavage from occurring.

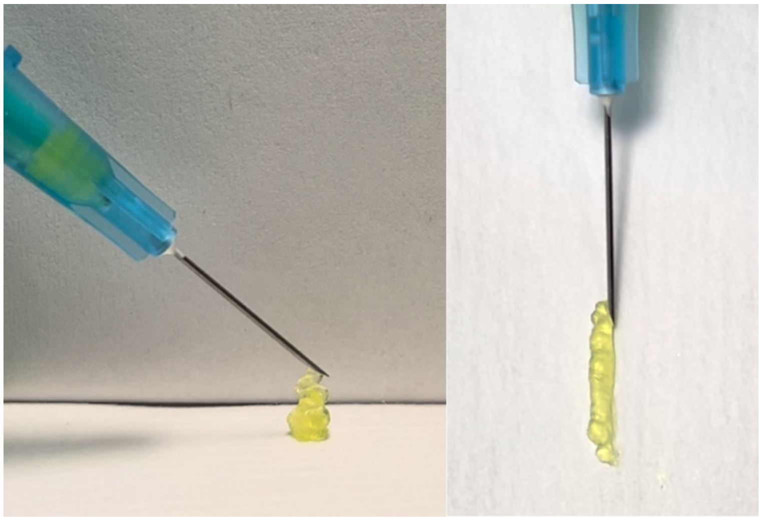

**Supplementary Fig. 3. Injection of PhoCoil gel.** 10 wt% gel was loaded into a 1 mL-syringe with a 25 gauge, 5/8 in needle. Injection required little pressure and the material retained gel-like properties post-injection as visualized.

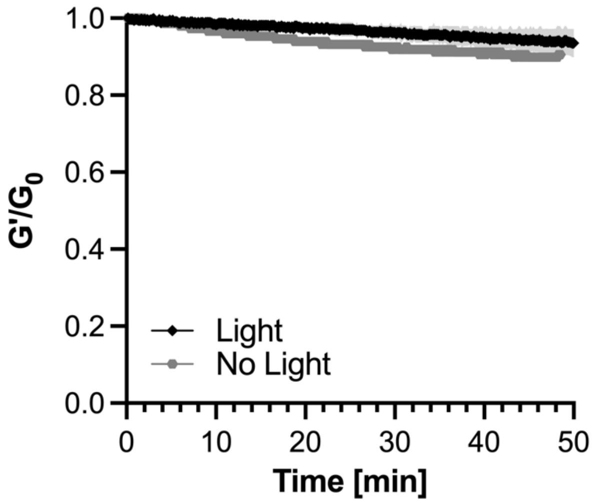


**Supplementary Fig. 4. Coil gel stiffness in response to light.** Coil gels, which are not designed to be light-responsive, were tested for potential photosoftening during an oscillatory rheology time sweep. Rheology was conducted at 5% strain and 10 rad $s^{-1}$ with 405nm light intensity set to 10 mW $cm^{-2}$ at the gel. $G'_0$ was defined as the storage modulus immediately before light exposure was initiated. 10 wt% gels were used, with three independent gels measured for the light condition, and one for the no light condition. Data is presented as the mean (line) ± SD (shaded area) for the light condition.

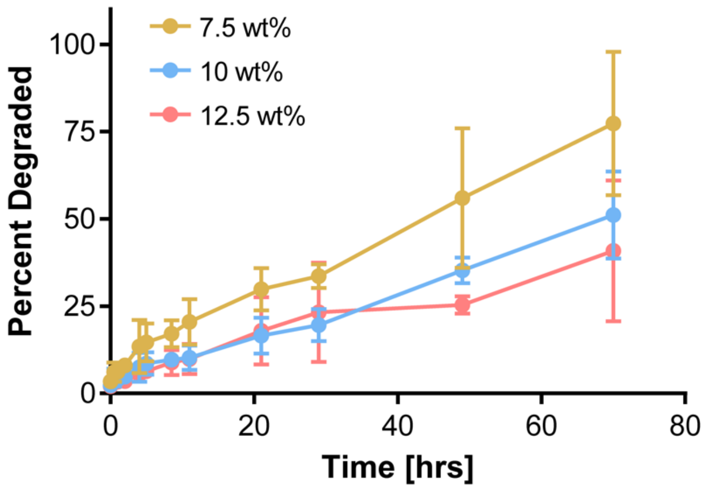


**Supplementary Fig. 5. Extended PhoCoil bulk gel degradation in ambient light.** 25 μL gels were covered with PBS and kept in ambient light. PBS was sampled at various intervals over 3 days and analyzed by BCA assay to determine protein content as a measure of the percentage of gel degraded at each time point. Three independent gels were tested for each weight percentage. Data shown are mean ± SD.

| Gel Weight Percentage (wt%) | 405 nm Light Exposure (min) | Half-life (hrs) |
|---|---|---|
| 7.5 | 0 | 44 ± 16 |
| | 30 | 6 ± 2 |
| | 60 | 2.2 ± 0.3 |
| | 90 | 2.4 ± 0.8 |
| 10 | 0 | 82 ± 32 |
| | 30 | 7.0 ± 0.6 |
| | 60 | 5.2 ± 0.5 |
| | 90 | 3 ± 1 |
| 12.5 | 0 | 112 ± 51 |
| | 30 | 9 ± 2 |
| | 60 | 5.1 ± 0.3 |
| | 90 | 4.0 ± 0.9 |

**Supplementary Table 1. PhoCoil bulk degradation half-lives.** Gel degradation curves in Figure 4B were used to calculate degradation half-lives for each condition. These values represent degradation of 25 μL gels covered with PBS in microcentrifuge tubes. Data shown are mean ± SD.

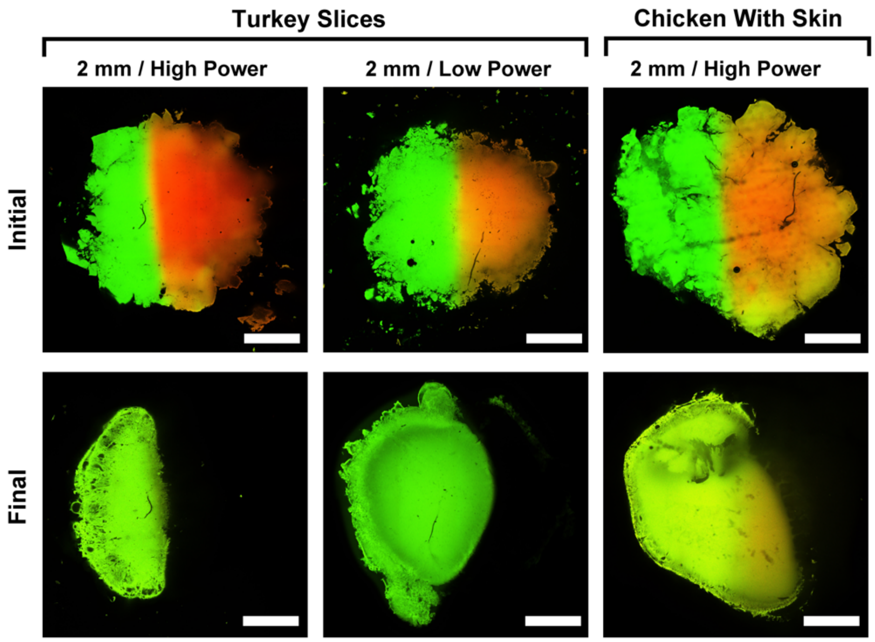


**Supplementary Fig. 6. Extended *ex vivo* photodegradation.** The left half of the gels was covered with a photomask, and the entire gel was placed under 2 mm of deli turkey or chicken with skin on. Gels were exposed to 405 nm light, for short periods at high power and long periods at low power, confocal imaged to visualize the pattern (initial), and left to degrade in excess PBS prior to final confocal imaging.
